# A neural network account of memory replay and knowledge consolidation

**DOI:** 10.1101/2021.05.25.445587

**Authors:** Daniel N. Barry, Bradley C. Love

## Abstract

Replay can consolidate memories through offline neural reactivation related to past experiences. Category knowledge is learned across multiple experiences, and its subsequent generalisation is promoted by consolidation and replay during rest and sleep. However, aspects of replay are difficult to determine from neuroimaging studies. We provided insights into category knowledge replay by simulating these processes in a neural network which approximated the roles of the human ventral visual stream and hippocampus. Generative replay, akin to imagining new category instances, facilitated generalisation to new experiences. Consolidation-related replay may therefore help to prepare us for the future as much as remember the past. Generative replay was more effective in later network layers functionally similar to the lateral occipital cortex than layers corresponding to early visual cortex, drawing a distinction between neural replay and its relevance to consolidation. Category replay was most beneficial for newly acquired knowledge, suggesting replay helps us adapt to changes in our environment. Finally, we present a novel mechanism for the observation that the brain selectively consolidates weaker information; a reinforcement learning process in which categories were replayed according to their contribution to network performance. This reinforces the idea of consolidation-related replay as an active rather than passive process.

## 1. Introduction

### 1.1 Memory consolidation-related replay

Memory replay refers to the reactivation of experience-dependent neural activity during resting periods. First observed in rodent hippocampal cells during sleep (Wilson and McNaughton 1994), the phenomenon has since been detected in humans during rest (Tambini and Davachi 2013; Hermans et al. 2017; Schapiro et al. 2018; Liu et al. 2019; Wittkuhn and Schuck 2021), and sleep (Schönauer et al. 2017; Zhang et al. 2018). These investigations have revealed replayed experiences are more likely to be subsequently remembered, therefore replay has been proposed to strengthen the associated neural connections and to protect memories from being forgotten. This memory consolidation-related replay can be viewed as distinct from task-related replay, the neural reactivation observed during task performance which supports cognitive processes such as memory recall (Jafarpour et al. 2014; Michelmann et al. 2019; Wimmer et al. 2020), visual understanding (Schwartenbeck et al. 2021), decision making (Liu et al. 2021), planning (Momennejad et al. 2018) and prediction (Ekman et al. 2017). While traditional perspectives view memory consolidation as a gradual process of fixation whereby memories are stabilised (Squire and Alvarez 1995; McGaugh 2000), in this paper we advocate the more contemporary view that offline consolidation-related replay is more dynamic in nature (Mattar and Daw 2018). Using a computational approach, we test hypotheses that offline replay may be a creative process to serve future goals, that it matters exactly where in the brain replay occurs, that it helps us at particular stages of learning, and that the brain might actively choose the optimal experiences to replay.

### 1.2 Generative replay of category knowledge

Neural replay which supports memory consolidation during rest and sleep has been traditionally assumed to be veridical, such that we commit the events of that day to long-term memory by replaying the episodes as they were originally experienced. However, there are circumstances in which this may be suboptimal or impractical. For example, a desirable outcome of category knowledge consolidation is to generalise to new experiences rather than recognise past instances. This phenomenon has been observed after sleep in infants (Gómez et al. 2006; Friedrich et al. 2015; Horváth et al. 2016), and in adults (Lau et al. 2011). Sleep also recovers the generalisation of phonological categories (Fenn et al. 2003), preserves generalisation performance in perceptual category learning (Graveline and Wamsley 2017), and assists in the abstraction of gist-like prototype representations (Lutz et al. 2017). It is still not understood how the brain consolidates and replays memory in the service of generalisation. In addition, although sleep benefits category learning for a limited number of well-controlled experimental stimuli (Schapiro et al. 2017), in the real world category learning takes place over many thousands of experiences, and storing each individual experience for replay is an impractical proposition. For these reasons, we propose the replay of novel, prototypical category instances would be a more efficient and effective solution. In fact, given the role of the hippocampus in both replay (Zhang et al. 2018) and the generation of prototypical concepts (Hassabis et al. 2007), we consider this the most likely form of category replay. While evidence for such generative replay of category knowledge has yet to be discovered in the human brain, replay of sequences immediately following task performance in humans has been shown to be flexible, in that items can be re-ordered based on previously learned rules (Liu et al. 2019). This is reminiscent of “pre-play” observed during task performance in rodents, where hippocampal “place cells” observed to fire in specific locations reactivate in a different order to represent a route which has not been taken before (Gupta et al. 2010).

Drawing inspiration from these observations, here we test the idea that replay which facilitates memory consolidation, occurring over extended offline time periods including sleep, might also be generative in nature, and that it’s flexibility may not just apply to the reorganisation of learned sequences, but the creation of entirely new instances of a category. While decoding the re-ordering of stimuli or route knowledge from brain data during replay has been shown to be a tractable approach, detecting entirely new instances of complex categories from the brain represents a significant challenge, and has yet to be demonstrated.

One approach to address this question is to simulate these processes in an artificial neural network. Prior research with artificial neural networks has modelled the replay of generated image stimuli (van de Ven et al. 2020). While revealing a promising avenue of investigation, the results of this study cannot be easily extrapolated to the brain or human visual experience. For example, the structure of only five convolutional layers in the network employed represents just a fraction of the size of larger models which have been shown to extract visual representations similar in nature to those processed by the brain (Schrimpf et al. 2018), whose complex structure can be compared to the ventral visual stream processing pathway, indicating a possible correspondence in functional architecture (Khaligh-Razavi and Kriegeskorte 2014; Güçlü and van Gerven 2015; Devereux et al. 2018), and whose object recognition performance approaches that of humans (He et al. 2015). Further, the networks employed by van de Ven et al. (2020) had limited visual experience, having been pre-trained on just 10 categories of objects. In contrast, an adult human brain will harbour a lifetime of visual knowledge which facilitates the learning of novel concepts. Therefore, to simulate the learning and generative replay of new categories realistically in adults, using an experienced network which contains a pre-existing vast body of knowledge about a range of other categories is an essential starting point. Another feature of the aforementioned study which limits the comparison to humans, is that the stimuli used were low-resolution photographs measuring 32 × 32 pixels, which do not reflect the complexity of human visual experience. To accurately simulate human learning and replay, much larger, high-resolution images which go some way towards approaching the complexity and richness of everyday human visual experience are required as training stimuli. Finally, prior attempts at replay in neural networks, whether generative (Kemker and Kanan 2017; van de Ven et al. 2020) or veridical (Hayes et al. 2021) have been deployed to address the “catastrophic forgetting” problem; the tendency of artificial networks to forget old categories when new ones are learned (Robins 1995; French 1999). While this has been proposed as a potential mechanism for why biological agents do not suffer from catastrophic forgetting, empirical evidence in support of this hypothesis has not been forthcoming to date. In addition, other solutions have been put forward on how brains and models may avoid catastrophic interference, such as Adaptive Resonance Theory (Grossberg 2013), and elastic weight consolidation (Kirkpatrick et al. 2017).

In this study, we investigated whether offline generative replay of novel concepts facilitated subsequent generalisation to new experiences using models which attempt to simulate the human brain and approximate more closely the visual environment in which it learns. To do this, we implemented generative replay in a well-studied deep convolutional neural network (DCNN), which consists of a complex architecture organised into five blocks of convolutional layers and boasts a high “brain-score” indicating the representations it extracts bear a similarity to those extracted by the brain and it performs favourably to humans in a categorisation task (Schrimpf et al. 2018). The network had prior experience of learning 1000 diverse categories of objects from over a million high-resolution complex naturalistic images, a process which is the network equivalent of a lifetime of visual experience and which yields within the model an optimised, high-functioning visual system. We tasked the model with learning 10 novel categories it had not seen before, using similarly high-resolution naturalistic images to those it has seen before, with an average resolution of around 400 × 350 pixels (Deng et al. 2009), representing an approximate 140-fold increase in visual details from stimuli used in prior work. The analogue in humans would be coming across 10 new categories we had not seen before and using our lifelong experience in processing visual information to extrapolate the relevant identifying features. After learning periods, we then simulated generative replay in the network, which attempted to mimic human consolidation during sleep, and monitored the network’s performance when it “woke up” the next day, to ascertain if we could provide computational support for the theory that such a process underlies the overnight improvements in generalisation observed in humans.

### 1.3 Effective neural loci of replay

Another outstanding question regarding replay, is despite being associated with subsequent memory (Zhang et al. 2018), it is not clear where in the brain replay makes a demonstrable contribution towards generalisation. Replay has been observed throughout the brain, early in the ventral visual stream (Ji and Wilson 2007; Deuker et al. 2013; Wittkuhn and Schuck 2021), in the ventral temporal cortex (Tambini et al. 2010; de Voogd et al. 2016), the medial temporal lobe (Staresina et al. 2013; Schapiro et al. 2018) the amygdala, (Girardeau et al. 2017; Hermans et al. 2017), motor cortex (Eichenlaub et al. 2020) and prefrontal cortex (Peyrache et al. 2009). It is not known if replay in lower-level brain regions actually contributes to the observed memory improvements or whether the key neural changes are made in more advanced areas, and this question cannot be answered using current neuroimaging approaches. One prior study has implemented replay within an artificial neural network from a single location at the end of the network (van de Ven et al. 2020). However, because the compact architecture of this network did not have a clear functional correspondence with information processing pathways in the brain, and because replay from other locations within the network was not also implemented for comparison, it is difficult to yield predictions from these results regarding effective replay locations in the human brain. In the current study, because we simulated replay in a neural network which processes images in a manner reflective of the human ventral visual stream, we could compare the effectiveness of replay from different layers with a purported representational correspondence to specific regions in the brain. In doing so, we aimed to make predictions about the effective cortical targets of offline memory consolidation in humans.

### 1.4 A time-dependent role for replay

Another open question regarding human replay is the duration of its involvement throughout the learning of novel concepts. It can take humans years to learn and consolidate semantic or conceptual knowledge (Manns et al. 2003), but neuroimaging studies of replay are limited to a time-span of a day or two, therefore it is still not known how long replay contributes to this process. Humans are thought to “reconsolidate” information every time it is retrieved (Dudai 2012), suggesting replay might play a continual role in the lifespan of memory. However recordings in rodents have shown that replay diminishes with repeated exposure to an environment over multiple days (Giri et al. 2019), suggesting the brain may only replay recently learned, vulnerable information. Answering this question in humans remains a challenge because of the impracticalities of tracking replay events for extended periods. Simulation in a human-like neural network represents a possible alternative to predict the relative contribution of replay to consolidation over long time-periods, an approach which has not been attempted to date. Here, we interleaved daily learning with nights of offline replay in a neural network which simulates the brain to understand at what stage in learning replay may be most effective in humans.

### 1.5 Replay of weakly-learned knowledge

An additional poorly understood principle of replay which we investigated in this study is why consolidation tends to selectively benefit weakly-learned over well-learned information (Kuriyama et al. 2004; Drosopoulos et al. 2007; McDevitt et al. 2015; Schapiro et al. 2018). Here, we modelled a candidate mechanism for how this occurs in the brain, by adding an auxiliary model (theoretically analogous to the hippocampus) to the neocortical-like model, which could autonomously learn the best consolidation strategy, determining what to replay and when.

### 1.6 Hypotheses

In addressing these outstanding questions regarding replay in the brain, we made a number of predictions. Because earlier brain regions are thought to extract equivalent basic features from all categories, we predicted replay of experience would be more effective in promoting learning at advanced stages of the network. We hypothesised the replay of “imagined” prototypical replay events would be as effective as veridical replay in helping us to generalise to new, unseen experiences, thus supporting our conceptualization of replay as a creative process. We predicted that the benefits of replay may be confined to early in the learning curve when novel category knowledge is being acquired. Finally, we hypothesised that a dynamic interaction between hippocampal and neocortical-like models would result in the prioritisation of weakly-learned items, in line with behavioural studies of memory consolidation.

## 2. Materials and Methods

### 2.1 Neural network

To simulate the learning of novel concepts in the brain, and test a number of hypotheses regarding replay, we trained a DCNN on 10 new categories of images. The neural network was VGG-16 (Simonyan and Zisserman 2014). This network is trained on a vast dataset of 1.3 million high-resolution complex naturalistic photographs known as the ImageNet database (Deng et al. 2009), which contains recognisable objects from 1000 categories in different contexts. The network learns to associate the visual features of an object with its category label, until it can recognise examples of that object which it has never seen before, simulating the human ability to generalise prior knowledge to new situations. The network takes a photograph’s pixels as input, and sequentially transforms this input into more abstract features. It learns to perform these transformations by adjusting 138,357,544 connection weights across many layers. Its convolutional architecture reduces the number of possible training weights by searching for informative features in any area of the photographs.

In these experiments, we task the VGG-16 network with learning 10 new categories of images. To do this, we retained the pre-trained “base” of this network, which consisted of 19 layers, organised into five convolutional blocks. Within each block there were convolutional layers and a pooling layer, with nonlinear activation functions. To this base, we attached two fully connected layers, each followed by a “dropout” layer, which randomly zeroed out 50% of units to prevent overfitting to the training set (Srivastava et al. 2014). At the end of the network a SoftMax layer was attached, which contained just 10 outputs rather than the original 1000, and predicted which of 10 classes an image belonged to. To facilitate the learning of 10 new classes, the weights of layers attached to the pre-trained base were randomly initialised. All model parameters were free to be trained. In total, 10 new models were trained, each learning 10 new and different classes.

### 2.2 Stimuli

Photographic stimuli for new classes were drawn randomly from the larger ImageNet 2011 fall database (Russakovsky et al. 2015), and were screened manually by the experimenter to exclude classes which bore a close resemblance to classes which VGG-16 was originally trained on. In total, 100 new classes were selected, and randomly assigned to the 10 different models to be trained. Within each class, a set of 1,170 training images, 130 validation images, and 50 test images were selected. The list of the selected classes is available in Supplementary Table 1.

### 2.3 Baseline training

We first trained a model without implementing replay, to serve as a baseline measure of network performance, and compare with other conditions which implemented replay. Ten models were trained on 10 new and different classes. To further prevent overfitting to the training set, images were augmented before each training epoch. This is similar to a human viewing an object at different locations, or from different angles, and facilitates the extraction of useful features rather than rote memorisation of experience. Augmentation could include up to 20-degree rotation, 20% vertical or horizontal shifting, 20% zoom, and horizontal flipping. Any blank portions of the image following augmentation were filled with a reflection of the existing image. Images were then pre-processed in accordance with Simonyan and Zisserman (2014). Depending on the experiment, the network was trained for 10 or 30 epochs. We used the Adam optimiser (Kingma and Ba 2014) with a learning rate of 0.0003. A small learning rate was chosen to reflect the fact that learning new categories in an adult human reflects a “fine-tuning” of an already highly-trained visual system. The training batch size was set to 36. The training objective was to minimise the categorical cross-entropy loss over the 10 classes. Training parameters were optimised based on validation set performance. We report the model’s performance metrics from the test set only. This is a collection of novel images from each category which the network does not learn nor is it tuned on, therefore reflecting the model’s ability to generalise to new stimuli after training, and is thus termed “generalisation performance” in the figures. Training was performed using TensorFlow version 2.2.

### 2.4 Replay

Replay was conducted between training epochs, to simulate “days” of learning and “nights” of offline consolidation. We conceptualised replay representations as generative, in other words they represented a prototype of that category never seen before, from a particular point in the network. To generate these representations, the network activations induced by the training images from the preceding epoch were extracted from a particular layer in the network using the Keract toolbox (Remy 2020). For each class separately, a multivariate distribution of activity was created from these activations using the SciPy toolbox (https://scipy.org/). This multivariate normal distribution is an extension of the one-dimensional normal distribution to higher dimensions, and is specified by its mean and covariance matrix. This resulted in a single unique distribution for each specific class, which represented the relationship between units of the layer which had been previously observed for that class. We then sampled randomly from this distribution, creating novel activation patterns for that class at that point in the network (Fig 1A). These novel activation patterns represented a prototype of that category. The end result was a representation that was a rough approximation of the layer’s representations of that category if a real image was processed, but novel in nature (see supplementary Fig 1). The human brain equivalent would be the approximate pattern of neural activity which is representative of that category at a particular stage in the ventral visual stream. In the brain, these hypothetical prototypical concepts would be likely generated from more high-level regions such as the hippocampus and prefrontal cortex (Hassabis et al. 2007; Bowman et al. 2020). Our model was generative as it could create new samples, however it offered several advantages over traditional generative models. We were not limited by a bottleneck symmetrical architecture, and our procedure allowed the model to learn generative samples at multiple levels of representation. Further, our model represented a proper vision model which showed parallels with the functional architecture of the ventral visual stream in the brain, whereas current generative models do not show this correspondence or scale well to such a deep architecture. Finally, our model is specialised for object recognition, with the resulting generated representations shaped by these task pressures.

**Fig 1.**
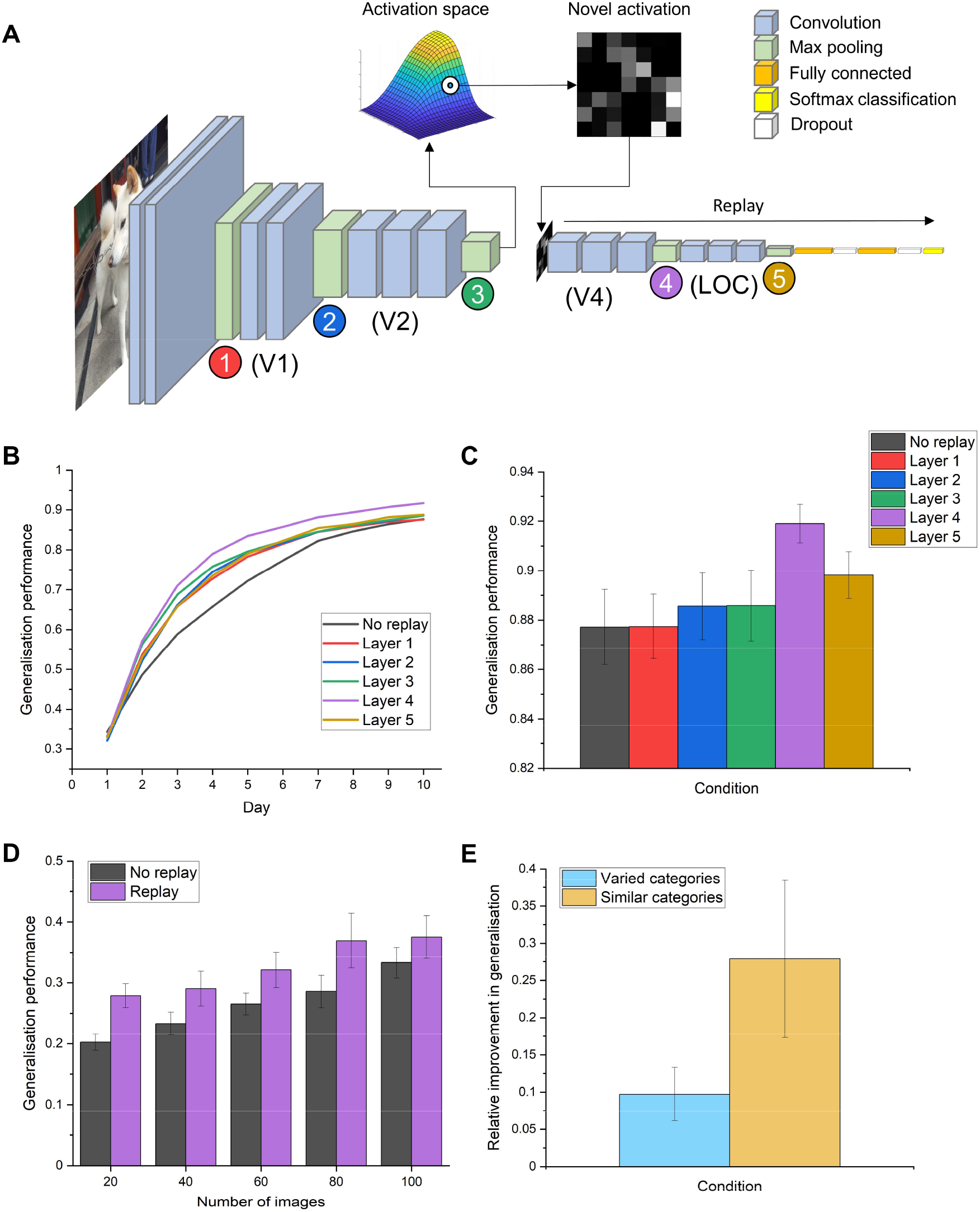
The effects of generative replay from different layers of a model of the human ventral visual stream on generalisation to new exemplars. (A) The VGG-16 network attempts to simulate the brain’s visual system by looking at photographs and extracting relevant features to help categorise the objects within. We trained this network on 10 new categories of objects it had not seen before. In between learning episodes, to simulate sleep-facilitated consolidation in humans, we implemented offline memory replay as a generative process. In other words, the network “imagined” new examples of a category based on the distribution of features it has learned so far for that object (activation space), and used these representations (novel representation) to consolidate its memory. The network did not create an actual visual stimulus to learn from, rather it recreated the neuronal pattern of activity that it would typically generate from viewing an object from that category. We display here an example of replaying from a mid-point in the network, but all five locations where replay was implemented are indicated by the coloured circles. The brain regions which have been reported to contain functionally similar representations to different network layers, derived from Güçlü and van Gerven (2015), are listed beneath. (B) The effects of memory replay from different layers on the network’s ability to generalise to new examples of the 10 categories, throughout the course of 10 learning episodes. Plotted values represent the mean accuracies from 10 different models which each learned 10 new and different categories. (C) The final recognition accuracies (+/- S.E.M.), averaged across 10 models, on the new set of photographs after 10 epochs of learning. We reveal the location in a model of the ventral stream where replay maximally enhances generalisation performance is an advanced layer which bears an approximate functional correspondence to the lateral occipital cortex (LOC) in humans. The benefits of replay from other locations were less pronounced, with the earliest layer showing the least benefit to generalisation. (D) The benefits of replay from layer four on generalisation performance with limited numbers of exemplars (E) The effect of generative replay from layer four on the generalisation performance of classes when learned alongside diverse categories or where all are conceptually similar.

The number of novel representations created for replay was equivalent to the number of original training images (1,170). To test where in the network replay is most effective, this process was performed at one of five different network locations, namely the max pooling layers at the end of each block (Fig 1A). For the first four pooling layers, creating a multivariate distribution from such a large number of units was computationally intractable, therefore activations for each filter in these layers were first down-sampled by a factor of eight for layer one, by four for layers two and three and two for layer four. The samples drawn from the resulting distribution were then up-sampled back to their original resolution. These lower-resolution samples are also theoretically relevant, in that they were created to mimic the schematic nature of mental and dream imagery which takes place during rest and sleep. To replay these samples through the network, the VGG-16 network was temporarily disconnected at the layer where replay was implemented, and a new input layer was attached which matched the dimensions of the replay representations. This truncated network was trained on the replay samples using the same parameters as regular training. We assume that the brain actively chooses to replay each concept learned that day, by reactivating the prototypical representations extracted from many experiences and the associated category label together during sleep. After each epoch of replay training, the replay section of the network was reattached to the original base, and training on real images through the whole network resumed. To assess the effects of generative replay on stimuli disambiguation, we took 10 classes from the 100 which were highly similar (plants, see supplementary table 2), and trained an additional network on these categories. We then assessed whether replaying similar classes in the same model led to a greater relative increase in class performance from baseline accuracies. We did this by dividing the increase in generalisation performance resulting from replay by the original baseline performance. To find out how many exemplars are needed for generative replay to have a beneficial effect on category learning we trained the same models with 20, 40, 60, 80 and 100 images, again for ten “days”, and replayed an equivalent number of generated representations in each case. To simulate veridical replay, in other words the replay of each individual experience as it happened, rather than the generation of new samples, we used the activations for each object at that layer in the network during replay periods. These were not down-sampled during the process. Given how many examples of a concept we generally encounter, veridical replay of all experience is not a realistic prospect, which is why prior attempts to simulate replay in smaller-scale networks have also avoided this scenario in their approaches (Kemker and Kanan 2017; van de Ven et al. 2020). To additionally demonstrate the improvements that replay affords on each day relative to the previous day, we calculated the performance improvement from day n to day n+1, divided by the difference between model performance on day n and 1, which represents the potential room for improvement.

### 2.5 Replay within a reinforcement learning framework

We tested a process through which items which are most beneficial for replay might be selected in the brain. We proposed that such selective replay may involve an interaction between the main concept learning network (VGG-16), and a smaller network which learned through reinforcement which concepts are most beneficial to replay through the main network during offline periods. The neural analogue of such a network could be thought of as the hippocampus, as the activity of this structure precedes the widespread reactivation of neural patterns observed during replay (Zhang et al. 2018). This approach is similar to the “teacher-student” meta-learning framework which has been shown to improve performance in deep neural networks (Fan et al. 2018). The side network was a simple regression network with 10 inputs, one for each class, and one output, which was the predicted value for replaying that class through the main network. Classes were chosen and replayed one at a time, with a batch size of 36. To train the side network, a value of 1 was inputted for the chosen class, with zeros for the others. The predicted reward for the side network was the change in performance of the main network after each replay instance, which was quantified by a change in chi-square; a contrast of the maximum number of possible correct predictions by the main network, versus its actual correct predictions. A positive reward was therefore a reduction in chi-square, which resulted in an increase in the side network’s weight for that class. This led to the class being more likely to be chosen in future, as the network’s weights were converted into a SoftMax layer, from which classes were selected probabilistically for replay. Through this iterative process, the side network learned which classes were more valuable to replay, and continually updated its preferences based on the performance of the main network. Reducing the chi-square in this dynamic manner improves the overall network accuracy as it progressively reduces the disparity between the network’s classifications and the actual class identities. To generate initial values for the side network, one batch of each class was replayed through the main network. The Adam optimiser was used with a learning rate of 0.001 and the objective was to minimise the mean squared error loss. The side network was trained for 50 epochs with each replay batch. The assessment of network improvement was always performed on the validation set, and the reported values are accuracy on the test set, reflecting the ability of the network to generalise to new situations.

## 3. Results

### 3.1 Localising where in the ventral visual stream generative replay is likely to enhance generalisation

We first sought to establish where in the visual brain the replay of category knowledge might be most effective in helping to generalise to new experiences, as the functional relevance of replay observed in many different brain regions has yet to be established. To obtain a baseline measure of how the network would perform without replay, the network learned 10 new categories in the absence of offline replay. Next, we implemented generative memory replay. To do this, we captured the “typical” activation of the network for a category and sampled from this gist-like representation to create novel, abstracted representations for replay (Fig 1A).

We simulated generative replay from different layers in the DCNN, equivalent to different brain regions along the ventral stream. Specifically, we trained the network over 10 epochs, mimicking 10 days of learning in humans, and replayed prototypical representations after each training epoch, simulating 10 nights of offline consolidation during sleep. In Fig 1B we show how replay affects the ability of the network to generalise to new exemplars of the categories over the course of learning. Replay substantially speeds up the learning process, with replay from layer four already reaching the final baseline generalisation performance three days earlier. Fig 1C shows the final best performing models in each replay condition. A one-way repeated-measures ANOVA on the final models revealed a difference across conditions (F_(5,45)_ = 7.23, p < 0.001), with planned Bonferroni-corrected post-hoc comparisons revealing that only replay from layer 4 (t_(9)_ = −4.31, p = 0.002) was significantly higher than baseline. We performed an additional analysis to confirm that the down-sampling of earlier layers did not explain this finding, by further down-sampling the replay representations in layer four by a factor of seven, and generalisation performance in this layer was still significantly higher than baseline (see supplementary Fig 2). Therefore, there is a differential benefit of replay throughout the network, where replay in the early layers is of limited benefit, whereas replay in the later layers boosts generalisation performance to a greater degree. This predicts that early visual areas in the brain may not store sufficiently complex category-specific representations, curtailing the effectiveness of generated replay representations, whereas areas further along the ventral visual stream, such as the lateral occipital cortex, might be better positioned to support the generation of novel, prototypical concepts which accelerates learning in the absence of real experience and helps us to generalise to new situations. We further investigated if generative replay could benefit category learning where few exemplars are available. In Figure 2D we show that generative replay from layer four could improve generalisation when learning and replaying just 20, 40 or 60 exemplars (all t-tests below Bonferroni-corrected threshold of p = 0.01). We also assessed the effects of replay on class disambiguation in this layer, by training a model containing conceptually highly similar classes collated from all of the other models, and comparing the relative increase in generalisation performance from the original class accuracies. Figure 2E shows a replay-induced performance increase for conceptually similar items, but this did not reach statistical significance (t_(9)_ = −2.10, p = 0.065).

**Fig 2.**
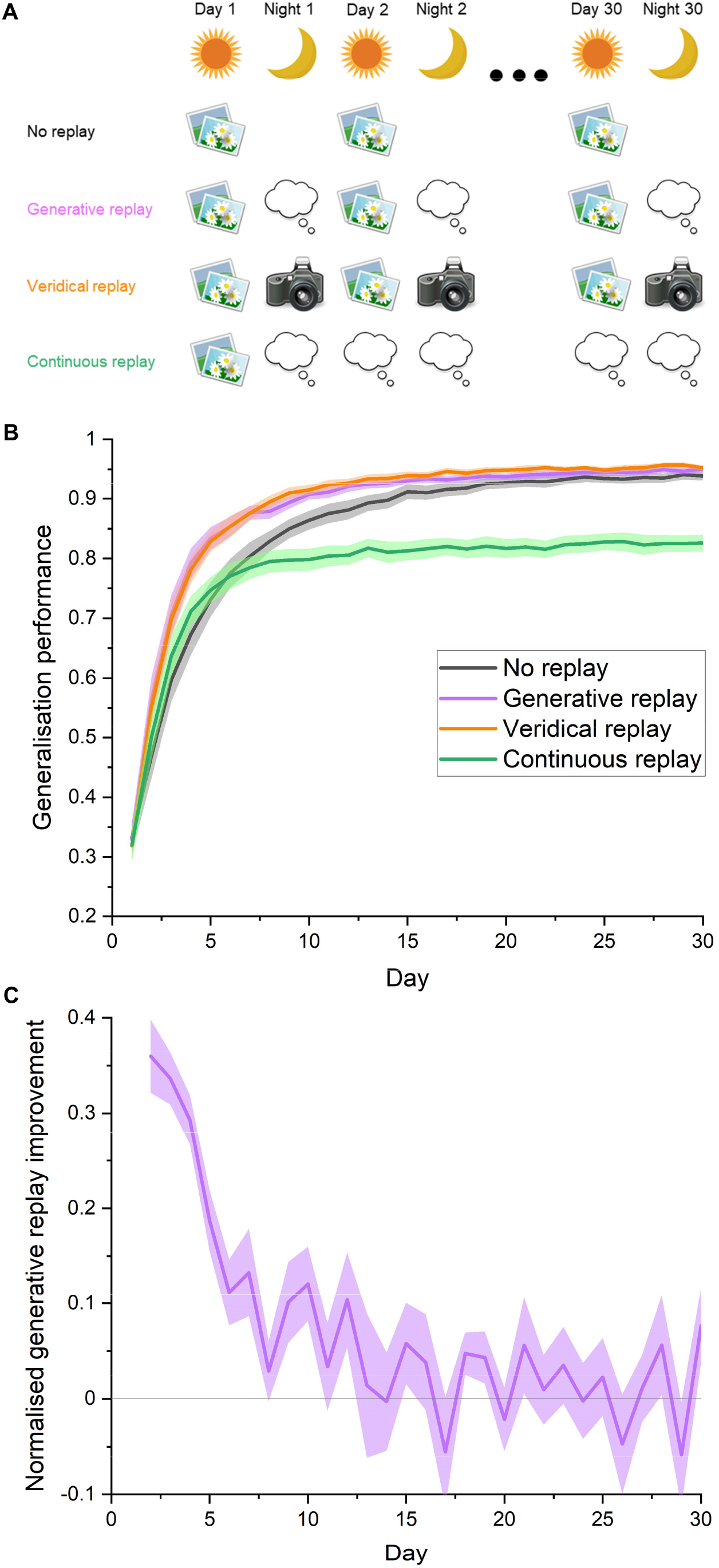
The facilitatory effects of memory replay across category learning. We simulate the long-term consolidation of category memory by extending training to 30 days. (A) Schematic showing the different experimental conditions. “No replay” involves the model of the visual system learning the 10 new categories without replay in between episodes. “Generative replay” simulates the brain imagining and replaying novel instances of a category during “night” periods of offline consolidation, from a layer bearing some functional approximation to the lateral occipital cortex. “Veridical replay” simulates the hypothetical performance of a human who, each night, replays every single event which has been experienced the preceding day. “Continuous replay” simulates a single day of learning, followed by days and nights of replay, investigating the potential benefit afforded by replay given only brief exposure to a category. For both day-time learning of real images and night-time consolidation of generated representations, the number of training stimuli was always 1,170 for each class. (B) The ability of the network to generalise to new exemplars of a category during each day throughout the learning process. Generalisation performance is measured by the proportion (+/- S.E.M.) of correctly recognised test images across 10 models. Generative replay maximally increases performance early in training, suggesting it might be optimal for new learning and recent memory consolidation. Despite being comprised of internally generated fictive experiences, generative replay was comparably effective to veridical replay throughout the learning process, positing it as an attractive, efficient and more realistic solution to memory consolidation which does not involve remembering all experiences. Continuous replay after just one day of learning substantially improved generalisation performance, but never reached the accuracy levels of networks which engaged in further learning. (C) The improvement in performance that generative replay affords on each day relative to the possible improvements from the previous day.

### 3.2 Tracking the benefits of replay across learning

In the second experiment, we extended training to 30 days of experience, interleaved with nights of offline generative replay to simulate learning over longer timescales and predict when in learning replay might be more effective (Fig 2A). Guided by the results of experiment one, we implemented replay from an advanced layer corresponding to the lateral occipital cortex. A mixed between-within ANOVA showed an interaction between condition and day (F_(29,522)_ = 5.03, p < 0.001) with planned post-hoc Bonferroni-corrected comparisons (p < 0.00167) revealing a difference between generative replay and baseline for days two to six, and eight (Fig 2B). Visualising the network’s improvement in performance from day to day relative to the potential room for improvement from the previous day confirmed that the benefits of generative replay were limited to early learning (Fig 2C). Therefore, offline generative replay might be more effective at improving generalisation to new exemplars at the earliest stages of learning. This suggests replay might facilitates rapid generalisation, which maximises performance given a limited set of experiences with a category.

We were interested to compare generative replay with the unlikely veridical, high-resolution scenario whereby humans could replay thousands of encounters with individual objects exactly as they were experienced. We termed this “veridical replay” (Fig 2A), which involved capturing the exact neural patterns associated with each experienced object during learning, and replaying these from the same point in the network. A mixed between-within ANOVA did not reveal any difference between generative and veridical replay in terms of generalisation performance (F_(1,18)_ = 0.16, p = 0.696), nor was an interaction effect observed between day and condition (F_(29,522)_ = 0.29, p = 0.999, Fig 2B). Therefore, generative replay was comparably effective to veridical replay of experience in consolidating memory, despite being entirely imagined from the networks prior experience. This provides tentative support for the hypothesis that generative replay is a putative form of category replay in humans, as it would appear vastly more efficient to imagine new concepts from an extracted prototype.

The aforementioned results simulated the benefits of replay under optimal conditions where humans encounter the same categories every day, however there are instances where exposure will be limited. To what extent can offline replay compensate for this limited learning? We simulated this in our model of the ventral stream by limiting the learning of actual category photographs to one day, and substituted all subsequent learning experiences with offline replay, termed “continuous replay” (Fig 2A). Despite the absence of further exposure to the actual objects, we found the network could increase its generalisation accuracy from 32% to 83% purely by replaying imagined instances of concepts it has partially learned. This result may inform our understanding of the human ability to quickly learn from limited experience. However, a mixed-between ANOVA revealed a statistically significant interaction effect between day and condition (F_(29,522)_ = 3.78, p < 0.001), with planned Bonferroni post-hoc comparisons revealing a difference between generative replay and continuous replay from day nine onwards (all p < 0.00167). Therefore, replayed representations appear to be dynamic in nature, as the prototypes generated from that first experience were not sufficient to train the network to its maximum performance, as is observed when learning and replay are interleaved. This suggests that replayed representations continue to improve as they are informed by ongoing learning, therefore generative replay in the human brain throughout learning may be envisaged as a constantly evolving “snapshot” of what has been learned so far about that category.

### 3.3 Determining how the brain might select experiences for replay

We proposed that replay may be a learning process in itself, whereby the hippocampus selects replay items, and learns through feedback from the neocortex the optimal ones to replay. In our previous simulations we selected all categories for replay in equal number, however to simulate the autonomous nature of replay selection in the brain, we supplemented our model of the ventral visual stream with a small reinforcement learning network, approximating the theoretical role of the hippocampus in deciding what to replay (Fig 3A). The hippocampus-like model could choose one of the 10 categories to replay, and received a reward from the main network for that action, based on the improvement in network performance.

**Fig 3.**
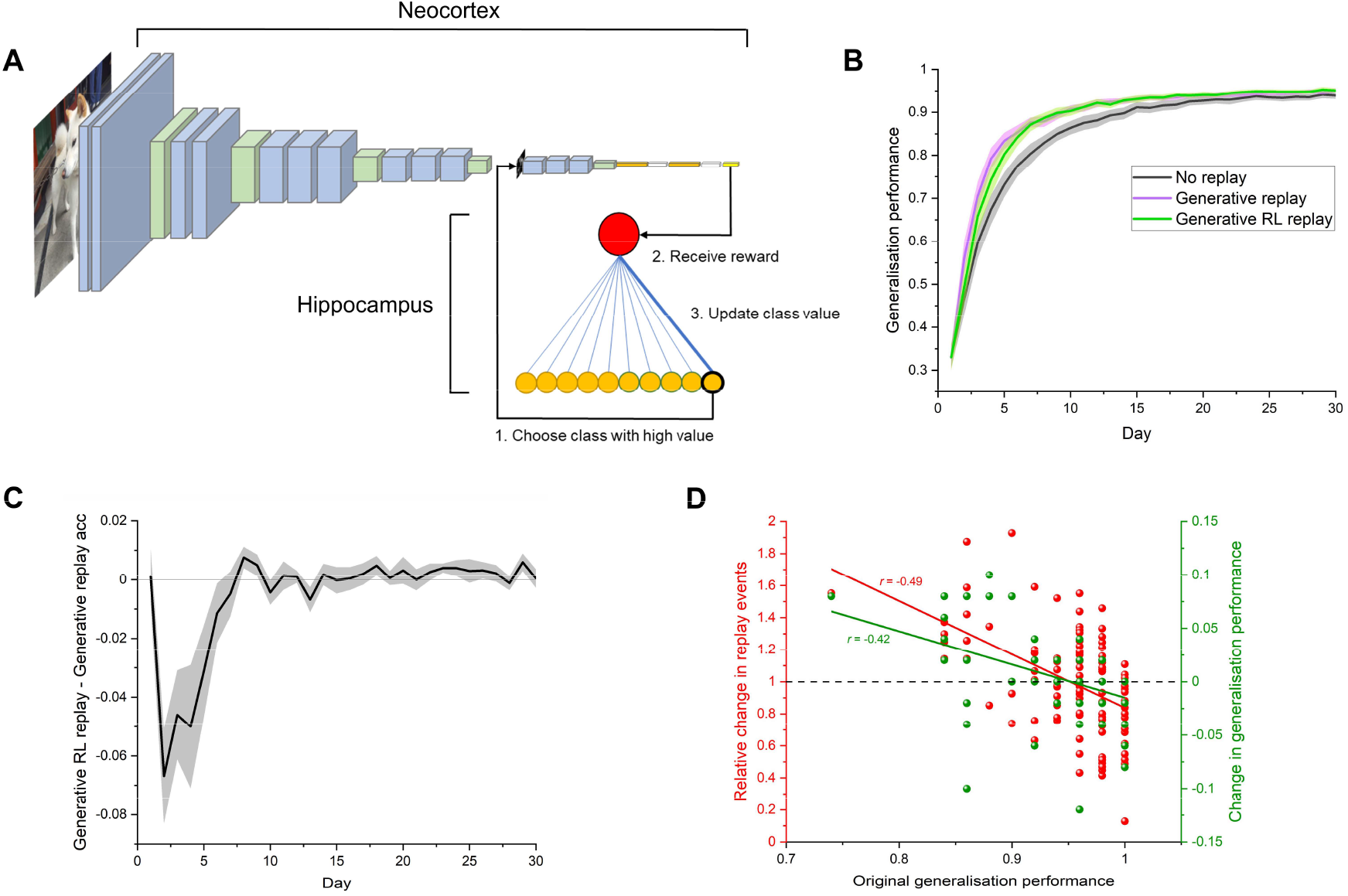
Replay as a reinforcement learning process simulates the brain’s tendency to consolidate weaker knowledge. (A) Replay in a model which approximates the visual system is controlled by a reinforcement learning (RL) network which aims to assume the role of the hippocampus. The RL network selects one of 10 categories to replay through the visual system and receives a reward based on the improved performance, learning through trial and error which categories to replay. (B) Overall generalisation performance on new category exemplars was similar for both generative replay and generative replay controlled by a reinforcement learning network. Generalisation performance represents mean accuracy (+/- S.E.M) on test images across 10 models which each learned 10 new categories. (C) The difference between generative replay and generative RL replay performance for each day. (D) The RL network learns to replay categories which were originally more difficult for the model of the visual system, and improves their accuracy. This effectively “rebalanced” memory such that category knowledge was more evenly distributed, and offers a candidate mechanism as to how the brain chooses weakly learned information for replay. Plotted values represent the 100 categories across 10 models. A proportion of the generalisation performance values are overlapping.

We trained our model of the visual system on 10 novel categories, implementing replay during offline periods as before, and compared its generalisation performance with that of the dual interactive hippocampal-cortical model. In terms of overall accuracy, although generative RL replay appeared to lag briefly behind generative replay at the beginning of training, both approaches performed similarly, with a mixed between-within ANOVA revealing no difference between the two conditions in terms of generalisation performance (F_(1,18)_ = 0.15, p = 0.704), nor was an interaction effect observed between day and condition (F_(29,522)_ = 1.28, p = 0.153, Fig 3B). Fig 3C plots the difference between the two conditions across learning. However, the reinforcement learning network which simulated the hippocampal replay systematically selected categories which were originally relatively weakly learned more often (R^2^ = 0.24, F_(1, 98)_ = 31.15, p < 0.001, Fig 3D), which resulted in their selective improvement (R^2^ = 0.18, F_(1, 98)_ = 21.15, p < 0.001). However, this came at a cost, with originally well-learned categories being replayed less often and a drop in their generalisation accuracy. We present the idea that such a reinforcement learning process may underlie the “rebalancing” of experience in the brain, and that replay may therefore help to compensate for the fact that some categories are more difficult to learn than others.

## 4. Discussion

We simulated the consolidation of category knowledge in a large-scale neural network model which approximates functional aspects of the human ventral visual system, by replaying prototypical representations thought to be formed and initiated by the hippocampus. The notion that replay of visual experiences might be generative in nature has been suggested by limited-capacity models which have been trained on low-resolution photographic images (van de Ven et al. 2020). However, our results using a model of the visual brain whose representations has compared favourably with actual brain data, represent more persuasive evidence that humans are unlikely to replay experiences verbatim during rest and sleep to improve category knowledge, and might be more likely to replay novel, imagined instances instead. In addition, the large number (117,000) of high-resolution complex naturalistic images we used for training in this experiment more closely reflected real-world learning and facilitated the extraction of gist-like features. While empirical evidence exists that humans replay novel sequences of stimuli (Liu et al. 2019), our work suggests that the brain might go further and uses learned features of objects to construct entirely fictive experiences to replay. We speculate that this creative process is particularly important for the consolidation of category knowledge as opposed to the replay of episodic memory (Deuker et al. 2013; Schapiro et al. 2018; Zhang et al. 2018), because of the requirement to abstract prototypical features and use these to generalise to new examples of a category. We propose that generative replay confers additional advantages such as constituting less of a burden on memory resources, as not all experiences need to be remembered. Further, our replay representations were highly effective in consolidating category knowledge despite being down-sampled, and these compressed, low-resolution samples would reduce storage requirements further. Perhaps the simulation that most favourably supported the hypothesis that category replay in the brain likely adopts this compressed, prototypical format is that it aided generalisation to a similar degree as the exact veridical replay of experience in boosting generalisation performance. Therefore, the main advantage to generative replay over veridical replay is that it represents a feasible, efficient solution to memory consolidation without compromising effectiveness. In addition, generative replay can add to events which have been experienced. Our findings therefore encourage a reconceptualization of the nature of consolidation-related replay in humans, that it is not only generative, but also low resolution or “blurry”, as is the case with internally generated imagery in humans (Giusberti et al. 1992; Lee et al. 2012). In fact, the kind of replay we propose here may be the driving force behind the transformation of memory into a more schematic, generalised form which preserves regularities across experiences while allowing unique elements of experience to fade (Love and Medin 1998; Winocur and Moscovitch 2011; Sweegers and Talamini 2014). The challenge for future empirical studies in humans to confirm our hypothesis, will be to decode prototypical replay representations during rest and sleep. In addition, future modelling and empirical work should address the sequential nature of learning and replay, as life experience does not consist of still snapshots of experience, such as those used in these experiments. Prior modelling work has shown that a video game-playing agent can improve its performance by learning inside its own generated environment (Ha and Schmidhuber 2018), which is more akin to an unfolding dream during sleep, and may provide inspiration for modelling the generative replay of video-like events to support category learning.

Simulating replay in a human-like network also allowed us to answer a question not currently tractable in neuroimaging studies: where in the visual stream is replay functionally relevant to consolidation? In a prior simulation of replay in a neural network, van de Ven et al. (2020) demonstrated generative replay could attenuate forgetting when performed after the final convolutional layer, but its effectiveness was not compared to earlier layers, and the network employed, consisting of five convolutional layers, had not been compared with the human visual system. Deeper networks, such as the one used here, consisting of 23 layers in total, organised into five blocks of convolutional layers, not only extract useful category features from naturalistic images, but representations in network layers have demonstrated a degree of representational correspondence with specific brain regions along the ventral visual stream (Khaligh-Razavi and Kriegeskorte 2014; Güçlü and van Gerven 2015; Devereux et al. 2018), albeit not capturing all observable variance (Xu and Vaziri-Pashkam 2021). In keeping with our observation that low-resolution, coarse, schematic replay was effective in helping the network to generalise, we found the most effective location for replay to be in the most advanced layers of the network, layers which are less granular in their representations. This region shares some functional similarities with the lateral occipital cortex in humans, a region which represents more complex, high-level features (Güçlü and van Gerven 2015). In contrast, generative replay from the earliest layers corresponding to early visual cortex was less effective. These layers are sensitive to low-level visual features such as contrast, edges and colour, therefore generating samples from these layers will yield rudimentary-level category-specific information, which are of limited utility for replay and generalisation. High-level representations on the other hand, may contain more unique combinations and abstractions of these lower-level features. We also found replay from the penultimate layer was more effective than the final layer, suggesting the optimal replay location represents a balance between the presence of sufficiently complex category information and the number of downstream neuronal weights available to be updated based on replaying these features. These findings may encourage a re-evaluation of the functional relevance of replay in early visual cortices in both animals and humans, and generate specific hypotheses for potential perturbation studies to investigate the effects of disruptive stimulation at different stages of the ventral stream during offline consolidation.

Our simulations also revealed a phenomenon never before tested in humans, that the effectiveness of replay depends on the stage of learning. We acquire factual information about the world sporadically over time across contexts, for example we may encounter a new species at a zoo one day, and subsequently see the same animal on a wildlife documentary, and so on. Ultimately the consolidation of semantic information in the neocortex can take up to years to complete (Manns et al. 2003). However, our simulations suggest that replay may be most beneficial during the initial encounters with a novel category, when we are still working out its identifiable features and have not yet learned to generalise perfectly to unseen instances. It is therefore possible humans replay a category less and less with increasing familiarity, and there is some support for this idea in the animal literature (Giri et al. 2019). We speculate that if this is the case, the enhanced effectiveness for recent memories may have an adaptive function, allowing us to generalise quickly with limited information. In fact, our simulations showed that after a single learning episode, replay can compensate substantially for an absence of subsequent experience. Our results provide novel hypotheses for human experiments, testing for an interaction between the stage of category learning and the extent of replay. The fact that replay early in the learning process was more effective provides further support for our proposal that vague, imprecise replay events are useful for generalisation, as the networks imaginary representations at that stage would be an imperfect approximation of the category in question. We acknowledge there may be a “ceiling effect”, whereby later in training there is no further room for improvement, however we would posit that over the human lifespan, we are operating in the non-converged portion of the learning curve that we display here.

Our results also represent the first mechanistic account of how the brain selects weakly-learned information for replay and consolidation (Kuriyama et al. 2004; Drosopoulos et al. 2007; McDevitt et al. 2015; Schapiro et al. 2018). The hippocampus triggers replay events in the neocortex (Zhang et al. 2018), with a loop of information back and forth between the two brain areas (Rothschild et al. 2017), although the content of this neural dialogue is not known. Our simulations suggest that the hippocampus may learn the optimal categories to replay based on feedback from the neocortex. Our results showed that such a process resulted in the “rebalancing” of experience in an artificial neural network, where generalisation performance was improved for weakly learned items, and attenuated for items which were strongly learned. A reorganisation of knowledge of this kind has been observed in electrophysiological investigations in rodents, where the neural representations of novel environments are strengthened through reactivation at the peak of the theta cycle, while those corresponding to familiar environments are weakened through replay during the trough (Poe et al. 2000). This more even distribution of knowledge could be adaptive in both ensuring adequate recognition performance across all categories and forming a more general foundation on top of which future conceptual knowledge can be built. There have been recent theoretical and empirical demonstrations of how items get selected for replay within a reinforcement learning framework, such as the “tagging” of items that elicit a large prediction error during the learning phase (Momennejad et al. 2018), and the replay of events that are more likely to be encountered in future and which lead to the highest reward (Mattar and Daw 2018; Liu et al. 2021). However, these accounts do not explain why even in the absence of such prediction errors, or without knowing the likelihood of future events, knowledge which has been weakly-learned during waking periods is consistently targeted for replay and consolidation during sleep (Kuriyama et al. 2004; Drosopoulos et al. 2007; McDevitt et al. 2015; Schapiro et al. 2018). Our interactive networks suggest that offline reinforcement learning could account for the selection of weakly-learned knowledge during the replay process itself, and future experiments could assess whether our models choose the same categories for replay as humans when trained on the same stimuli.

In summary, our simulations provide supportive evidence that category replay in humans is a generative process and make the prediction that it is functionally relevant at advanced stages of the ventral stream. We have generated hypotheses about when during learning replay is likely to be effective and offer a novel account of replay as a learning process in and of itself between the hippocampus and neocortex. We hope these findings encourage a closer dialogue between theoretical models and empirical experiments. These findings also add credence to the emerging perspective that deep learning networks are powerful tools which are becoming increasingly well-positioned to resolve challenging neuroscientific questions (Richards et al. 2019).

## Acknowledgements

We thank the Love Lab for helpful discussions on this project. This work was supported by the Royal Society (grant number 183029) and the Wellcome Trust (grant number WT106931MA) to B.C.L. The funders had no role in study design, data collection and analysis, decision to publish, or preparation of the manuscript.

## Author Contributions

D.N.B: Conceptualization, methodology, software, data curation, investigation, formal analysis, visualization, writing-original draft preparation, writing-review & editing. B.C.L.: Conceptualization, methodology, resources, funding acquisition, supervision, writing-review & editing.

## Data and Code Availability

The code, environment, and additional information required to run the simulations is available at https://github.com/danielbarry1/replay.git and in the supplementary information. All relevant data in the paper is available at https://doi.org/10.6084/m9.figshare.14208470.

---

**Supplementary Fig 1.**
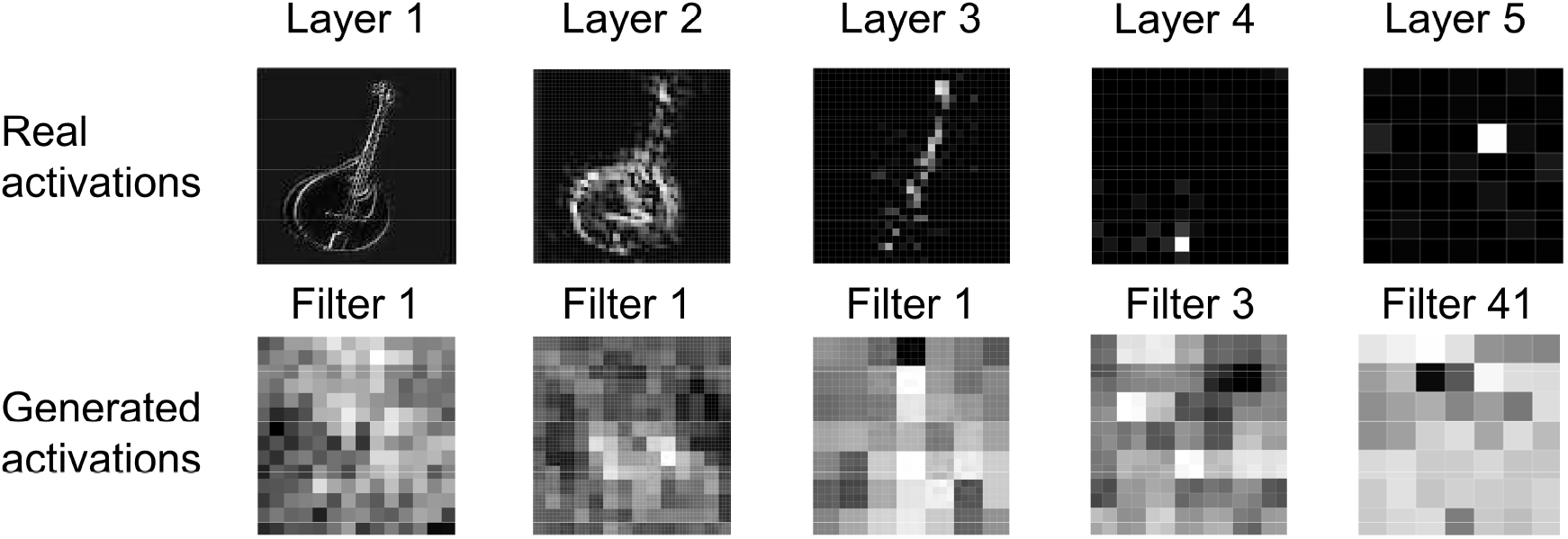
Samples of real and generated activations from different layers in the network, displayed as greyscale images. The first filters where information was visible is displayed.

**Supplementary Fig 2.**
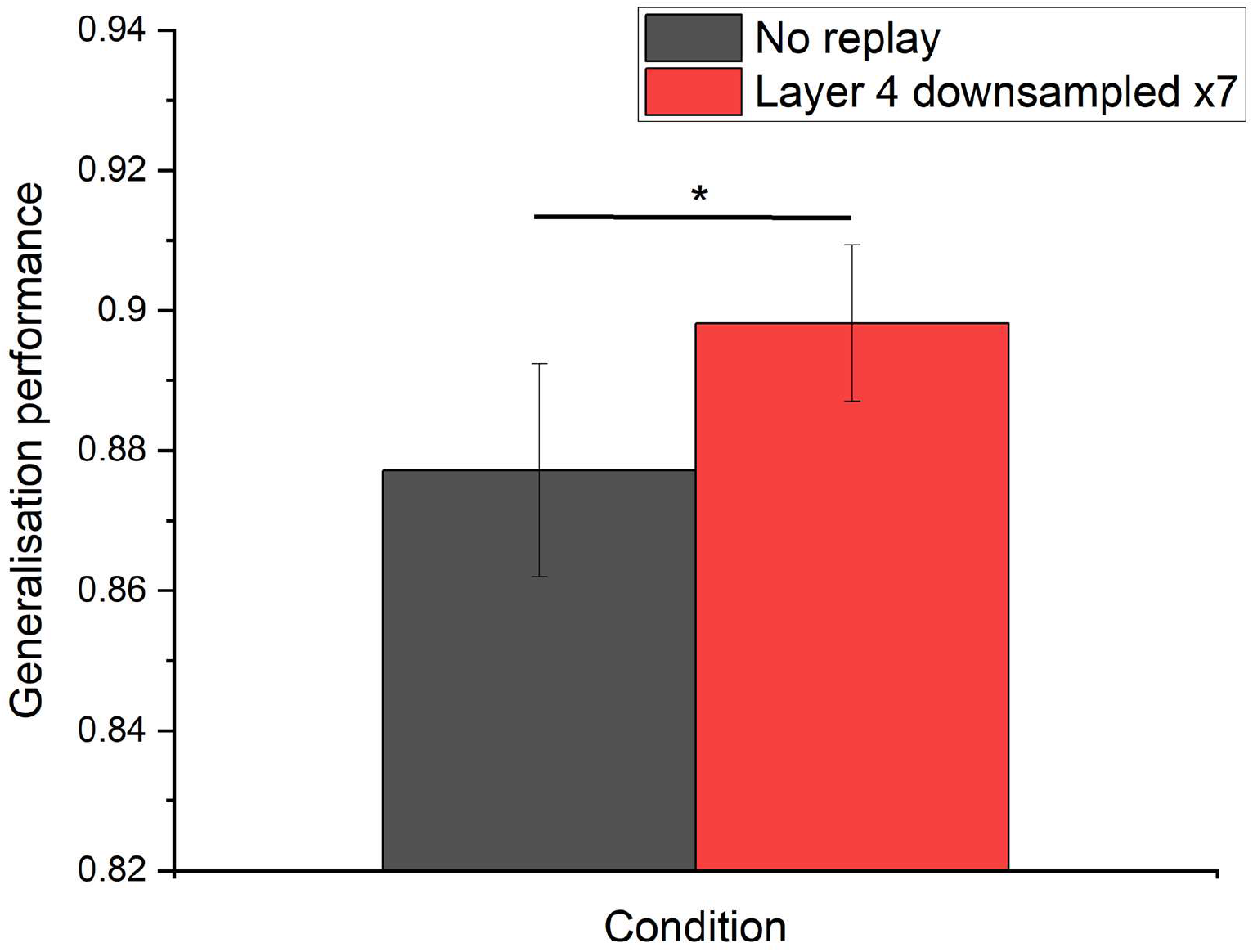
Generalisation performance improvement over baseline for replay from layer four when downsampled by a factor of seven.

**Supplementary table 1:**
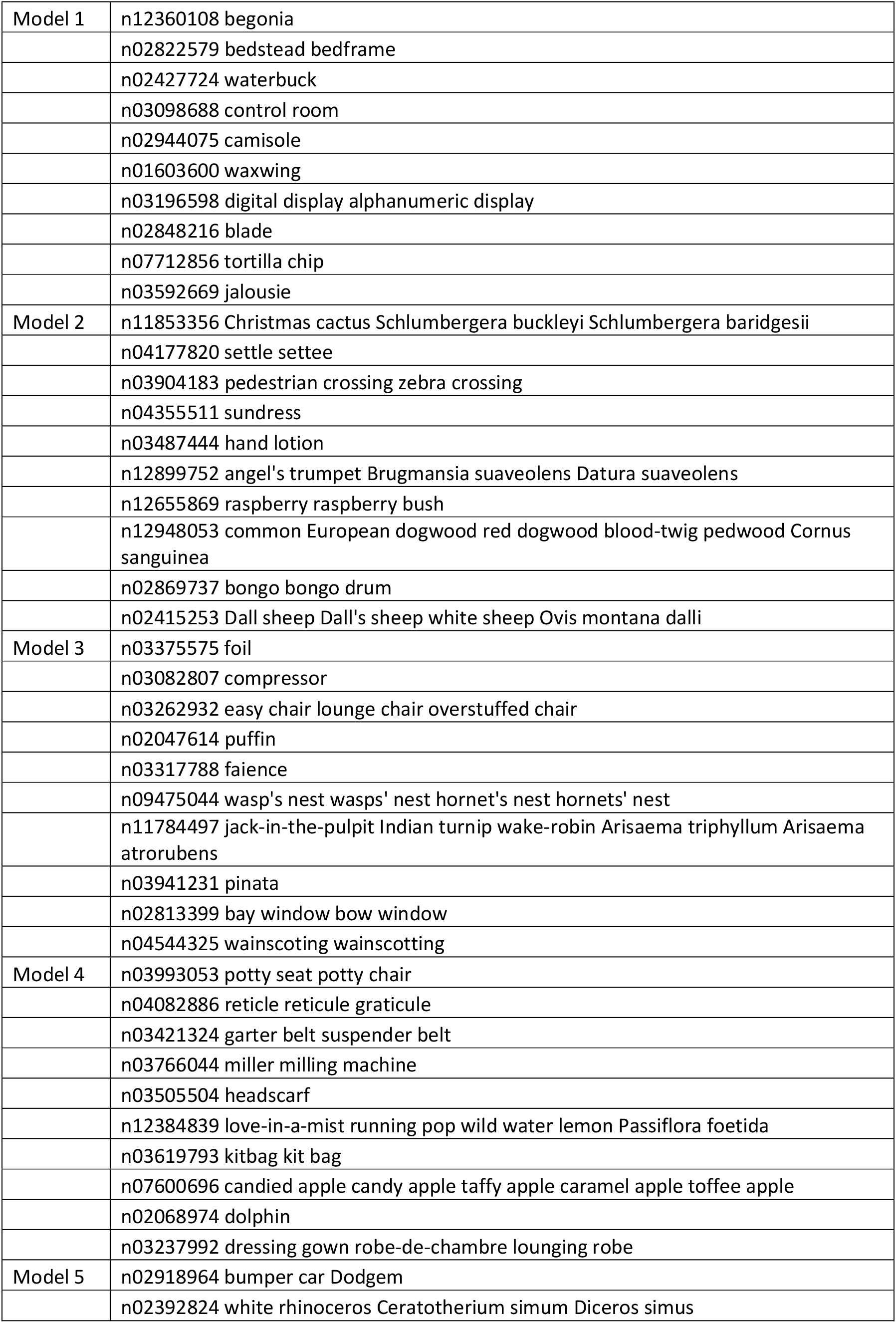
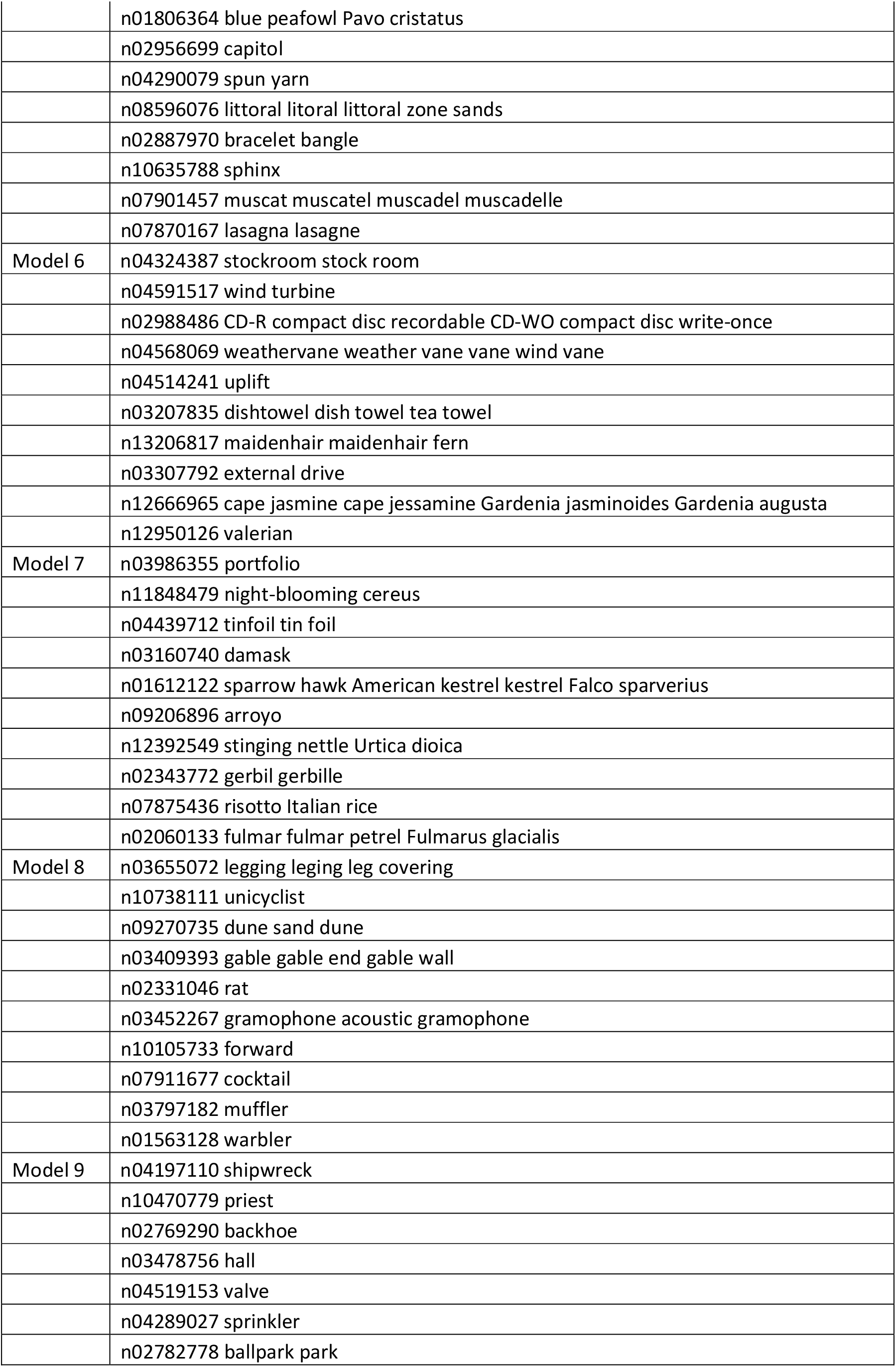
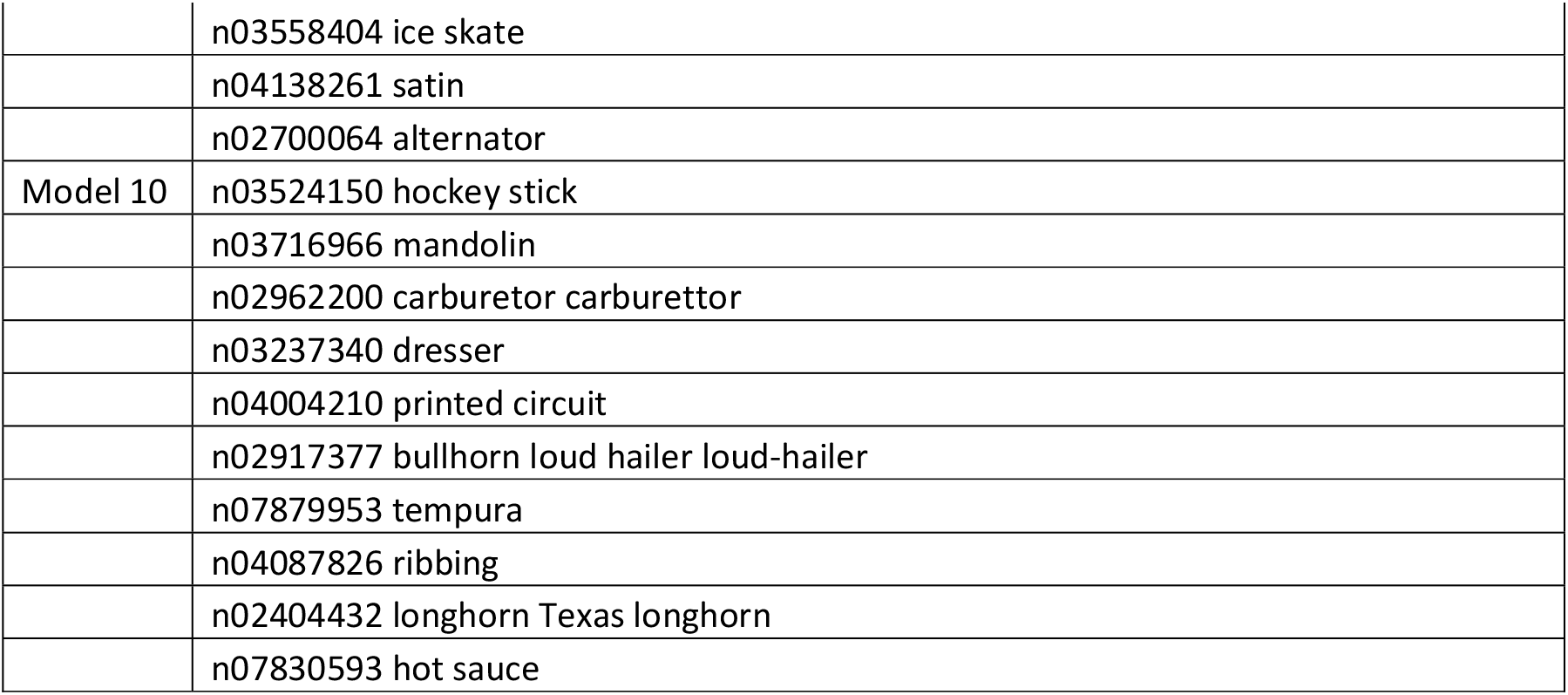
List of ImageNet classes by model

**Supplementary table 2:**
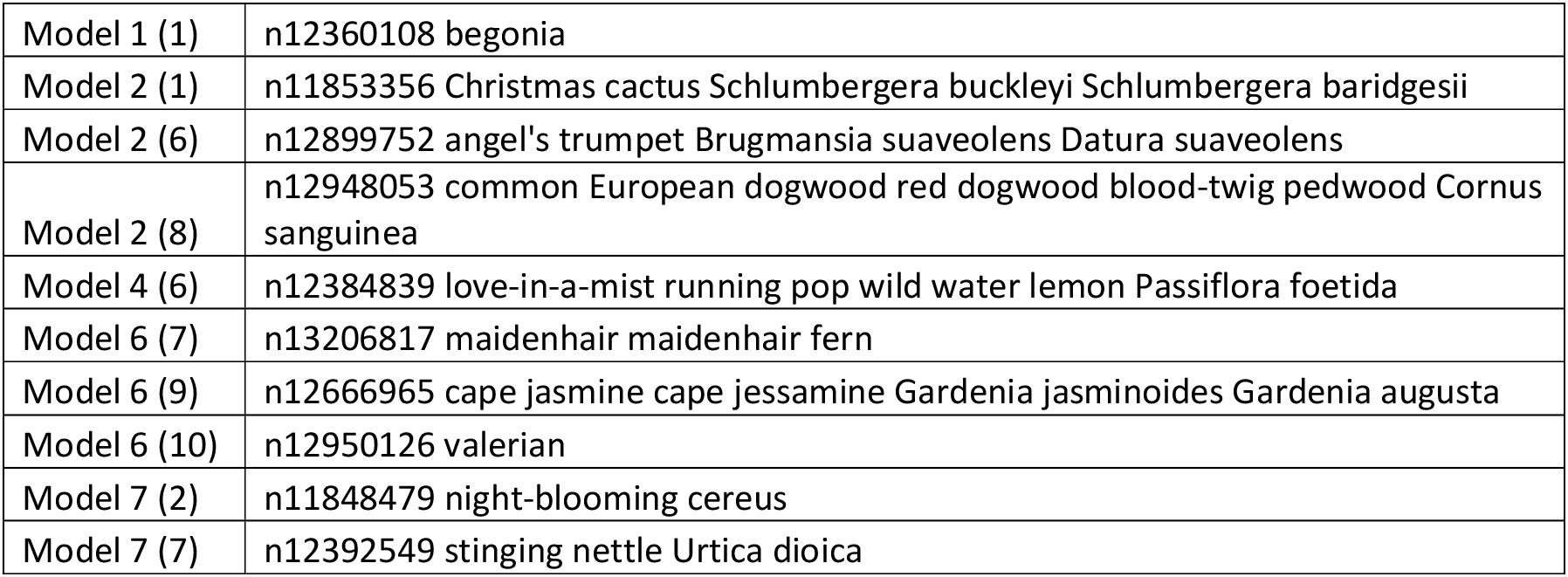
Conceptually similar classes (plants)

